# Off target: herbicides applied in cereal fields exclude non-competitive species, while replacing them by competitive weeds

**DOI:** 10.1101/2025.05.21.654858

**Authors:** Ségolène Humann-Guilleminot, Vincent Bretagnolle, Sabrina Gaba

## Abstract

Weed diversity plays an important role in maintaining resilient agroecosystems, yet agricultural practices, such as pesticide applications, significantly shape weed communities. Previous studies have primarily focused on comparing organic and conventional farming, with a particular emphasis on land-use intensification, with less attention on the impact of pesticides. Our study examined the impact of herbicide, fungicide, and insecticide use – both in terms of Treatment Frequency Index (TFI), i.e. intensity of application, and quantity – on weed communities in 96 non-organic cereal fields over 4 years. Interestingly, our results indicate that fungicide application intensity decreased weed abundance by 11.6% and species richness by 14.6% for each one-standard-deviation (1-SD) increase in TFI, whereas the total quantity applied (QA) did not. In contrast, for herbicides, QA had a stronger negative impact on weed communities than TFI, with a 14.3% decrease in weed abundance and a 12.4% decrease in species richness for each 1-SD increase in QA. This suggests that TFI of fungicide, which may reflect low-dose but frequent applications, could exert indirect and long-term effects on weed suppression, potentially by altering symbiotic arbuscular mycorrhizal fungi in the soil. In comparison, QA of herbicide more directly reflects the toxic load delivered to weeds and is therefore a better predictor of their suppression. Based on a combination of multivariate and univariate analyses and the study of underlying mechanisms in community changes, our results reveal that herbicides are the main factor shaping weed communities by decreasing the abundance of non-target/non-competitive species while replacing them by problematic, competitive to crops, weed species. These findings point out the potential ineffectiveness of herbicides application on problematic weeds and emphasize the need to reconsider the use of herbicides to maintain weed diversity in agricultural landscapes.

## 1. Introduction

Weeds play a critical ecological role within agroecosystems, by providing habitats and food for wildlife, supporting plant-pollinator interactions, enhancing soil health, facilitating nitrogen fixation, and preventing erosion (Altieri, 1999; Bretagnolle and Gaba, 2015; Marshall et al., 2003; Storkey et al., 2011). Unlike natural ecosystems, agroecosystems face frequent and severe disturbances such as pesticide use, soil tilling, and crop harvesting, which significantly impact their dynamics (Duru et al., 2015; Gaba et al., 2014; Tilman et al., 2002). Herbicides, along with mechanical weeding and tillage, are key methods used to manage weeds and minimize crop competition to protect yields by ∼20%, although the relationship between herbicide use and yield is not straightforward (Benaragama et al., 2016; Das et al., 2024; Gaba et al., 2014; Oerke, 2006; Tilman et al., 2002; Zimdahl, 2018). This frequent disturbance alters the plant community’s typical structure and diversity, often leading to fields dominated by one or two weed species (e.g. *Alopecurus myosuroides* or *Avena fatua* in cereals crops; Borgy et al., 2012; Storkey and Neve, 2018). Consequently, weed populations develop unique structures and traits influenced by both farming-induced disturbances and competition with these dominant crops (Gaba et al., 2014; Mahaut et al., 2020).

Numerous studies have compared how organic and conventional farming practices influence weed communities, with organic farming, which exclude synthetic pesticides and nitrogen inputs, generally promoting greater weed species richness (e.g., Bengtsson et al., 2005; Carrié et al., 2022; Gong et al., 2022; Henckel et al., 2015). However, organic farming also involves more diverse crop rotations and semi-natural habitats, which contribute to increased weed diversity (Tuck et al., 2014). This complexity makes it difficult to attribute differences in weed diversity solely to pesticide use. Other research has categorized the impacts of disturbances such as herbicides, nitrogen inputs, and tillage on weed abundance and diversity (Campiglia et al., 2018; Fonderflick et al., 2020; Fried et al., 2008; José-María et al., 2010; Storkey et al., 2011). At the species level, herbicides reduce weed abundance and species richness, with potential impact on non-target plants (Andert et al., 2022; Boutin et al., 2014).

However, at the community level, we still ignore whether community shifts resulting from herbicide use, are gradual or threshold-based, an important distinction for anticipating ecosystem responses to anthropogenic pressures and providing effective management strategies (Hillebrand et al., 2020; Spake et al., 2022). This raises questions about the role of herbicides as environmental filters that select for herbicide-resistant weed species (e.g., blackgrass and ryegrass) that might disrupt key biotic interactions, such as competition, resulting in species turnover or nested changes along an herbicide gradient. Additionally, nitrogen fertilization eliminates slow-growing species, and when combined with herbicides, further selects for nitrophilous and herbicide-tolerant weeds, reducing overall diversity (Berquer et al., 2023; Fonderflick et al., 2020). Furthermore, studying the gradients of fungicides and insecticides alongside herbicides offers a more holistic perspective on how pesticide use affects weed communities both directly (herbicides) or indirectly (other pesticides) by, for example, altering soil microbiome or reduce pollination.

In this study, we focused on the effect of pesticides on weed communities in 96 conventional winter cereal fields (2017-2020) located within a Long-Term Socio-Ecological Research platform in West of France. Because organic farming systems involve multiple, distinctly different management practices that may introduce additional variability than the sole exclusive use of pesticides, we chose to focus our study exclusively on non-organic cereal fields to isolate and quantify the specific effects of pesticide on weed communities. Based on a combination of multivariate and univariate analyses, our aim was to address four key questions: 1) How do herbicide, fungicide, and insecticide inputs influence weed communities at different scales, from species level to communities? 2) Do changes in weed community response to herbicide use occur gradually, or do they shift abruptly once a certain herbicide application threshold is reached? 3) What are the underlying mechanisms of community change, particularly in terms of nestedness and turnover components along the pesticide gradient? 4) Which species exhibit key pesticide-driven patterns in weed community structure, revealing both the species most susceptible to pesticide use and those that thrive under increased application intensity? We conducted all analyses using both the Treatment Frequency Index, which reflects an intensity of use, and the quantity of active ingredient applied. Although not independent of one another, the two indices represent different aspects of pesticides use allowing us to contrast the effects of disturbances related to pesticides use. Notably, we predicted that higher intensity of application would lead to gradual changes in weed species richness and abundance by consistently disrupting recovery, whereas higher quantity of active ingredient would cause more abrupt shifts due to its stronger direct impact on weed survival.

## 2. Results

We recorded a total of 181 weed species over the four years. Descriptive statistics (median, mean ± standard deviation, minimum and maximum) are presented in Table S1. This level of diversity is consistent with previous studies in European arable systems, where typical weed species richness in cereal fields ranges from 40 to 150 species depending on landscape complexity, management intensity, and sampling scale (e.g., Andert et al., 2022; Gaba et al., 2010; Hyvönen et al., 2003; José-María et al., 2010).

### 2.1. Effect of pesticides and nitrogen on weed abundance and species richness

The linear mixed-effect model (LMM) with the Treatment Frequency index (i.e. intensity of application; TFI) as the dependent variable indicate no statistical evidence that herbicides influence weed abundance but evidence that it decreases weed species richness (p = 0.10 and p = 0.09, respectively, effect size *r* = -0.18 for weed species richness; Table 1). However, we found that fungicides influence both weed abundance and species richness (p = 0.01 and *r* = -0.27 for both variables; Table 1, Figure 1). By contrast, no effect of insecticides is detected (p > 0.45; Table 1), though their use on wheat in our study area is rather marginal. Weak statistical evidence suggests that the quantity of active ingredient (QA) of herbicides influence weed abundance (p = 0.05 and *r* = -0.21), while it negatively impacts weed species richness (p = 0.02 and *r* = -0.24, respectively; Table 1; Figure 1). Additionally, nitrogen input increased weed abundance (p = 0.02, r = 0.25; Table 1) but not weed species richness (TFI and QA; p= 0.1 and p = 0.05, respectively; Table 1). Lastly, the sampling date did not influence weed abundance nor species richness.

**Figure 1:**
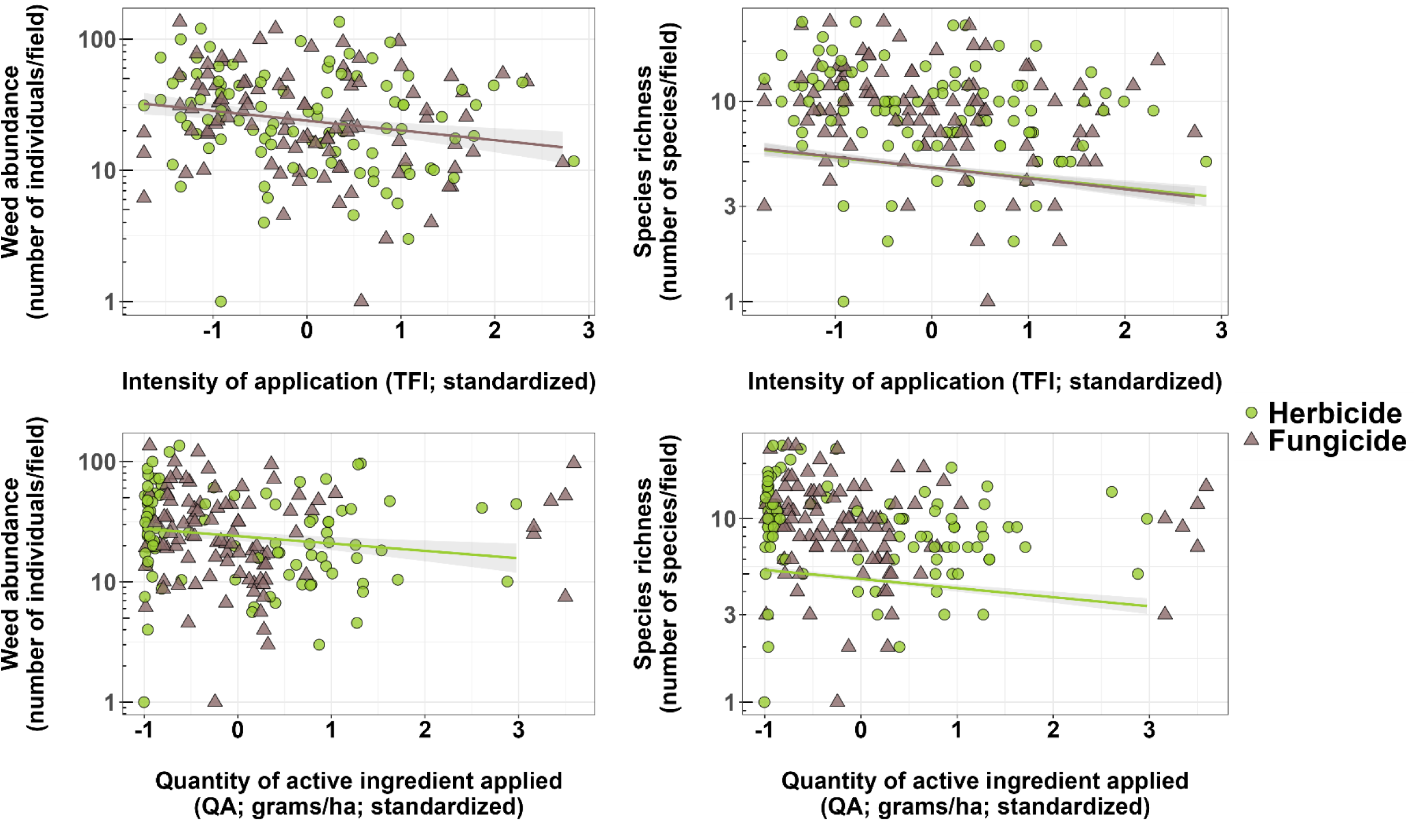
Weed abundance and species richness in relation to intensity (TFI) and quantity (QA) of herbicides (represented by green dots) and fungicides (represented by brown triangles). Both TFI and QA values were centered and scaled (i.e., standardized). A change of one unit (x-axis) corresponds to an increase or a decrease of one standard deviation from the mean of TFI or QA of herbicides or fungicides. Linear regression lines are added based on the statistical evidence: no linear regression is displayed in the absence of evidence of correlation, while one or two linear regression lines are presented when there are evidence supporting a relationship between one or more variables. Grey area represents the 95% confidence interval for the fitted line derived from predictions of the linear mixed-effects model, accounting for the effects of other variables in the model. A log scale is used on the y-axis to reflect that the response variable was log-transformed in the model.

**Table 1:** Summary of the linear mixed model investigating the effect of Treatment Frequency Index and Quantity of Active Ingredient of herbicide, fungicide, insecticide, nitrogen input and Julian date (all variables standardized) on log-transformed weed abundance and log-transformed species richness in field centres using a restricted maximum likelihood function. The table shows model estimates ± standard error (Est. ± s.e.) and associated 95% confidence intervals (C.I.). The table also shows multiplicative estimates for untransformed variable (Est. mult.) and associated multiplicative confidence intervals (C.I. mult.). Additionally, effect sizes *r* are provided, along with p-values that are derived from a t-test testing against the null hypothesis that the estimate is 0.

| Explanatory variables | Log-transformed weed abundance |  |  |  |  |  | Log-transformed species richness |  |  |  |  |  |
| --- | --- | --- | --- | --- | --- | --- | --- | --- | --- | --- | --- | --- |
|  | Treatment Frequency Index |  |  |  |  |  |  |  |  |  |  |  |
|  | Estimate ± s.e. | C.I. | Est. Mult. | C.I. mult. | r | p | Estimate ± s.e. | C.I. | Est. Mult. | C.I. mult. | r | p |
| Intercept | 3.13 ± 0.28 | [2.58 – 3.68] | 22.85 | [13.16 – 39.68] | - | < 0.001 | 2.16 ± 0.1 | [1.96 – 2.35] | 8.65 | [7.11 – 10.51] | - | <0.001 |
| Herbicide | -0.13 ± 0.08 | [-0.28 – 0.03] | 0.88 | [0.76 – 1.03] | -0.17 | 0.1 | -0.09 ± 0.05 | [-0.20 – 0.01] | 0.91 | [0.82 – 1.01] | -0.18 | 0.087 |
| Fungicide | -0.22 ± 0.08 | [-0.39 – -0.06] | 0.8 | [0.68 – 0.95] | -0.27 | 0.01 | -0.16 ± 0.06 | [-0.28 – -0.04] | 0.85 | [0.76 – 0.96] | -0.27 | 0.01 |
| Insecticide | -0.00 ± 0.08 | [-0.16 – 0.15] | 1 | [0.86 – 1.16] | -0.01 | 0.95 | -0.04 ± 0.06 | [-0.15 – 0.07] | 0.96 | [0.86 – 1.07] | -0.08 | 0.455 |
| Nitrogen input | 0.21 ± 0.09 | [0.04 – 0.39] | 1.24 | [1.04 – 1.48] | 0.25 | 0.02 | 0.1 ± 0.06 | [-0.02 – 0.23] | 1.11 | [0.98 – 1.26] | 0.18 | 0.1 |
| Julian date | -0.07 ± 0.09 | [-0.25 – 0.12] | 0.93 | [0.78 – 1.12] | -0.08 | 0.45 | -0.02 ± 0.06 | [-0.14 – 0.11] | 0.98 | [0.87 – 1.12] | -0.03 | 0.8 |
| Quantity of Active Ingredient |  |  |  |  |  |  |  |  |  |  |  |  |
|  | Log-transformed weed abundance |  |  |  |  |  | Log-transformed species richness |  |  |  |  |  |
|  | Estimate ± s.e. | C.I. | Est. Mult. | C.I. mult. | r | p | Estimate ± s.e. | C.I. | Est. Mult. | C.I. mult. | r | p |
| Intercept | 3.14 ± 0.27 | [2.60 – 3.69] | 23.1 | [13.39 – 39.85] | - | < 0.001 | 2.16 ± 0.09 | [1.99 – 2.34] | 8.71 | [7.29 – 10.42] | - | <0.001 |
| Herbicide | -0.15 ± 0.08 | [-0.30 – -0.00] | 0.86 | [0.74 – 1.00] | -0.21 | 0.05 | -0.13 ± 0.06 | [-0.24 – -0.02] | 0.88 | [0.79 – 0.98] | -0.24 | 0.02 |
| Fungicide | -0.07 ± 0.09 | [-0.25 – 0.10] | 0.93 | [0.78 – 1.10] | -0.09 | 0.39 | -0.1 ± 0.06 | [-0.23 – 0.02] | 0.90 | [0.79 – 1.02] | -0.17 | 0.1 |
| Insecticide | 0.10 ± 0.08 | [-0.05 – 0.26] | 1.11 | [0.95 – 1.30] | 0.14 | 0.19 | 0.04 ± 0.06 | [-0.07 – 0.15] | 1.04 | [0.93 – 1.17] | 0.07 | 0.49 |
| Nitrogen input | 0.23 ± 0.09 | [0.04 – 0.42] | 1.26 | [1.05 – 1.52] | 0.25 | 0.02 | 0.13 ± 0.07 | [0.00 – 0.26] | 1.14 | [1.00 – 1.30] | 0.23 | 0.05 |
| Julian date | -0.05 ± 0.10 | [-0.25 – 0.15] | 0.95 | [0.78 – 1.16] | -0.05 | 0.62 | 0.02 ± 0.07 | [-0.11 – 0.16] | 1.02 | [0.89 – 1.17] | 0.04 | 0.76 |

In field margins, none of the pesticides either measured by TFI or QA impact weed abundance and species richness. However, there is statistical evidence of a positive effect of nitrogen addition on weed species richness (Table S2; supporting information).

### 2.2. Weed community response to pesticide use

The multivariate analysis using distance-based redundancy analysis (dbRDA) provides strong statistical evidence that variation in weed community composition across winter cereal fields is explained by both TFI and QA of herbicides, and nitrogen input (Table 2, Figure 2), with a stronger effect of herbicide compared to nitrogen and from QA compared to TFI of herbicide (Table 2, Figure S1; supporting information). The model using QA explain slightly more variance in community composition (7.08%) than the TFI model (6.52%). In both models, the first dbRDA axis account for most of the variance, mainly driven by herbicides and nitrogen inputs (Table 2, Figure S1). Herbicide impact on weed communities is stronger than that of nitrogen input in both models (see effect sizes; Table 2). There is also a wide variation in weed communities across fields, with a gradual change in communities as herbicide use (both TFI and QA) increase (Figure 2).

**Figure 2:**
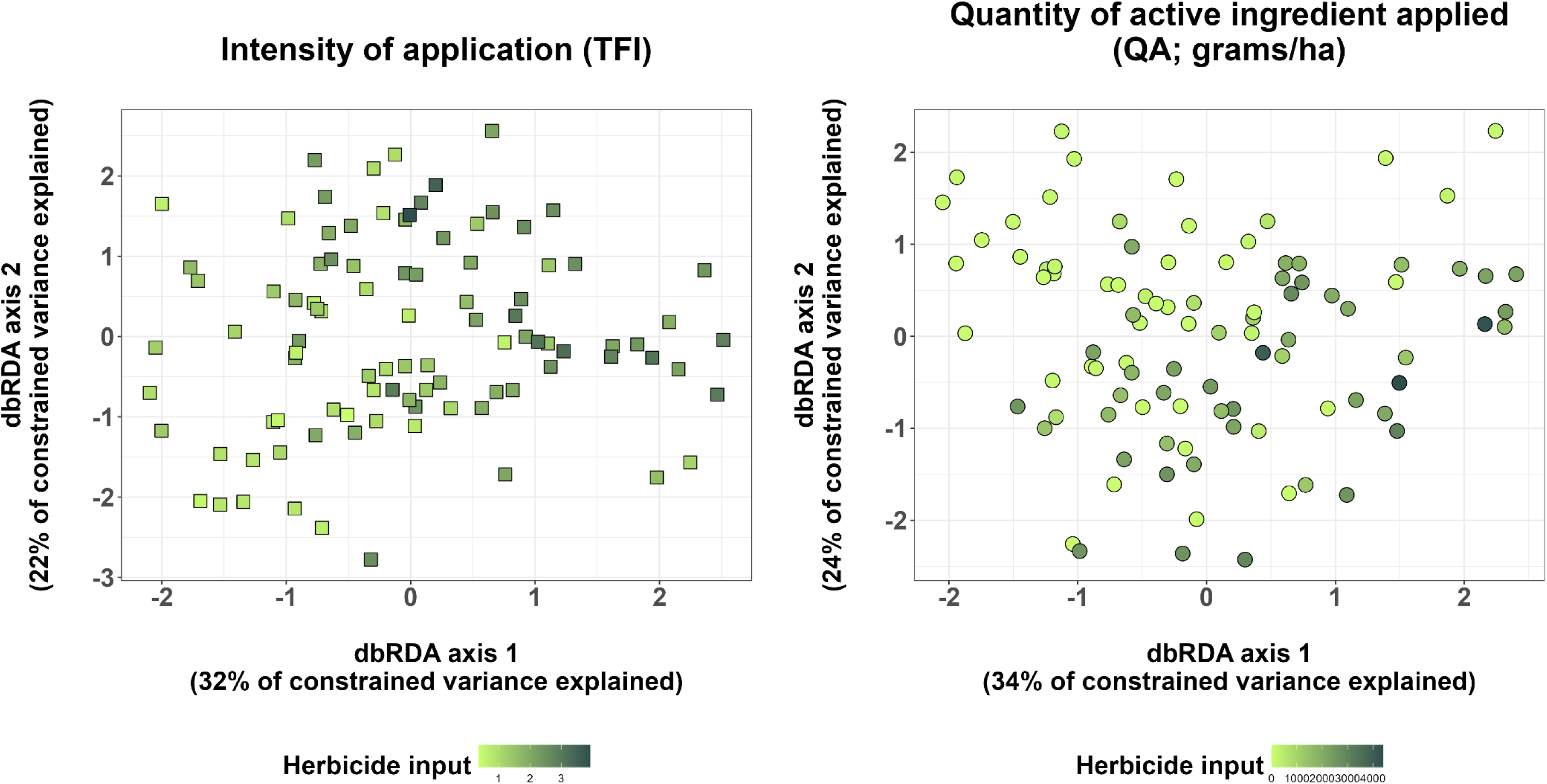
dbRDA biplot projections displaying weed community composition along herbicide input gradients based on the intensity of application of herbicide (Treatment Frequency Index, TFI; left) and the quantity of active ingredient applied (QA; right). Each point represents a community from a cereal field, with the colour gradient (light to dark green) indicating increasing herbicide input. In both models, herbicide input contributes most strongly to the first dbRDA axis (dbRDA1). The models explain 6.52% (TFI) and 7.08% (QA) of the variance in community composition.

**Table 2:** Summary statistics from the dbRDA model and ANOVA using Canberra distance to test the influence of TFI/QAs of herbicide, fungicide, insecticide, nitrogen inputs, and Julian date on weed community composition in field centres. The table shows the total contribution of each variable to the variance in community composition and the proportion of variance explained by each axis. It also includes the coefficients for each variable on the ordination axes, representing their coordinates along each axe. Additionally, effect sizes, indicating the strength of the influence of each variable on the ordination, are provided, along with p-values. Effects sizes correspond to the length of each arrow in the biplot and are calculated as the Euclidean distance from the origin to the variable’s coordinates on dbRDA1 and dbRDA2 (Figure S1). All variables were standardized (centered and scaled).

| Treatment Frequency Index (total contribution of all variables = 7.08%) |  |  |  |  |
| --- | --- | --- | --- | --- |
| Explanatory variables | Coefficient associated with each explanatory variable on each axis |  | Effect size | p |
|  | dbRDA1 | dbRDA2 |  |  |
| Herbicide | 0.74 | 0.46 | 1.20 | 0.001 |
| Fungicide | -0.14 | 0.55 | 1.04 | 0.11 |
| Insecticide | -0.17 | 0.41 | 0.28 | 0.10 |
| Nitrogen input | -0.48 | 0.02 | 0.87 | 0.009 |
| Julian date | -0.28 | 0.56 | 1.07 | 0.06 |
| Proportion explained of each axis: |  |  |  |  |
| dbRDA 1 = 32.31% (2.29% of variance explained by dbRDA1 overall) |  |  |  |  |
| dbRDA 2 = 22.12% (1.57% of variance explained by dbRDA2 overall) |  |  |  |  |
| Quantity of Active ingredient (total contribution of all variables = 6.52%) |  |  |  |  |
| Explanatory variables | Coefficient associated with each explanatory variable on each axis |  | Effect size | p |
|  | dbRDA1 | dbRDA2 |  |  |
| Herbicide | -0.60 | -0.47 | 1.36 | 0.005 |
| Fungicide | 0.04 | -0.66 | 0.87 | 0.24 |
| Insecticide | 0.14 | -0.10 | 0.67 | 0.74 |
| Nitrogen input | 0.51 | -0.20 | 0.74 | 0.003 |
| Julian date | 0.16 | -0.66 | 0.98 | 0.11 |
| Proportion explained of each axis: |  |  |  |  |
| dbRDA 1 = 34.44% (2.24% of variance explained by dbRDA1 overall) |  |  |  |  |
| dbRDA 2 = 24.25% (1.57% of variance explained by dbRDA2 overall) |  |  |  |  |

To investigate which species were most affected by herbicide use through the dbRDA analysis (TFI or QA) and contributed to shaping weed communities, we extracted and projected species scores along the two dbRDA axes. Because herbicide use was the main driver of community changes in both models, we reoriented the dbRDA reference frame (represented by red arrows in Figure S1) to align it with the herbicide vectors (TFI and QA), allowing a clearer projection of species scores along these gradients. After identifying the species most strongly associated with herbicide gradients, we searched a posteriori different sources (e.g., scientific literature and agricultural reference sites https://wikiagri.fr/) to determine whether these species are generally considered competitive or non-competitive in cereal cropping systems – regardless of their specific association with high or low herbicide input. The species classified as competitive or non-competitive based on agronomic knowledge are highlighted in the table associated to the Figure 3. A few species – *Mercurialis annua*, *Polygonum aviculare*, *Atriplex patula*, *Chenopodium album* and *Veronica persica* (later named non-competitive species; Figure 3) are associated with low herbicide input (either TFI or QA of herbicide). In contrast, *Lolium multiflorum, Galium aparine*, *Alopecurus myosuroides, Bromus hordeaceus,* and *Fallopia convolvulus* (later named as competitive species; Figure 3) are associated with higher herbicide use (Figure 3, Figure S1).

**Figure 3:**
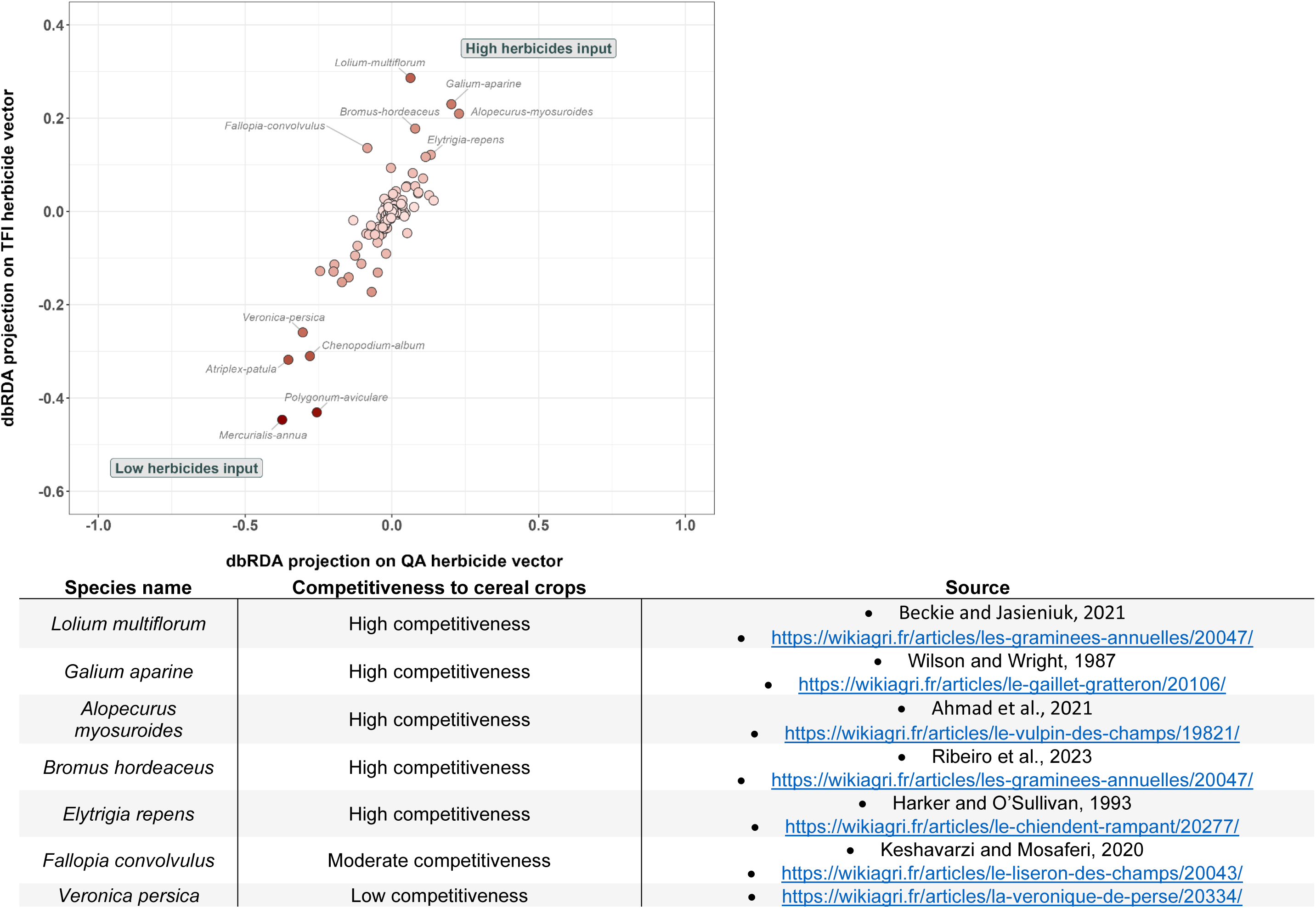
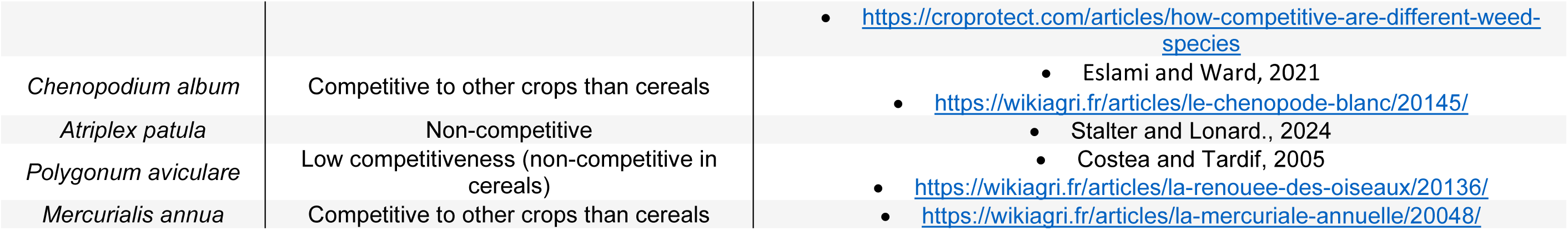
Species associations with the Treatment Frequency Index (TFI) and Quantity of Active Ingredient (QA) of herbicides represented by projecting species scores onto the two herbicide input vectors from the distance-based redundancy analysis (dbRDA). We reoriented the orthonormal reference frame (shown in Figure S1) to use the TFI and QA herbicide vectors as reference axes for projecting species scores associated with these metrics. Species associated with low herbicide input appear in the bottom left (either TFI or QA), while those associated with high herbicide input are positioned in the top right. The red colour gradient indicates the degree of association with the herbicide input vectors, with deeper red reflecting a stronger association with high or low herbicide sensitivity. This table accompanies figure 3 and classifies species which are at the two ends of the herbicide input gradient, i.e., the species that weigh the most in the analysis, as either competitive or non-competitive in cereal cropping systems. This classification was done independently of their scores on the gradient.

In field margins, neither pesticide use measured as TFI nor nitrogen input affect weed communities. However, there is weak statistical evidence suggesting that the QA of herbicides has an impact on weed communities. Additionally, the sampling date influence weed communities in both models (Table S3, Figure S2; supporting information).

### 2.3. Impact of pesticides on abundance of competitive and non-competitive weed species

Our analysis provides strong statistical evidence that both the Treatment Frequency Index (TFI) and the Quantity of Active Ingredient (QA) of herbicides affect the abundance of non-competitive to crops weed species (both p < 0.001 and *r =* -0.41 and *r =* -0.44, respectively). Conversely, we found no impact of herbicide input on the abundance of competitive weeds (p = 0.62 and p = 0.99, for TFI and QA, respectively; Table 3, Figure 4). Additionally, while nitrogen input is found to influence the abundance of non-competitive species, it has no effect on competitive species (Table 3). This influence of nitrogen is comparatively weaker than that of herbicides, with effect sizes of ***r*** *=* 0.29 and ***r*** *=* 0.37 in TFI and QA models, respectively. We report no statistical evidence that any other variables, including fungicides and insecticides, affect the abundance of any weed species subgroup (Table 3).

**Figure 4:**
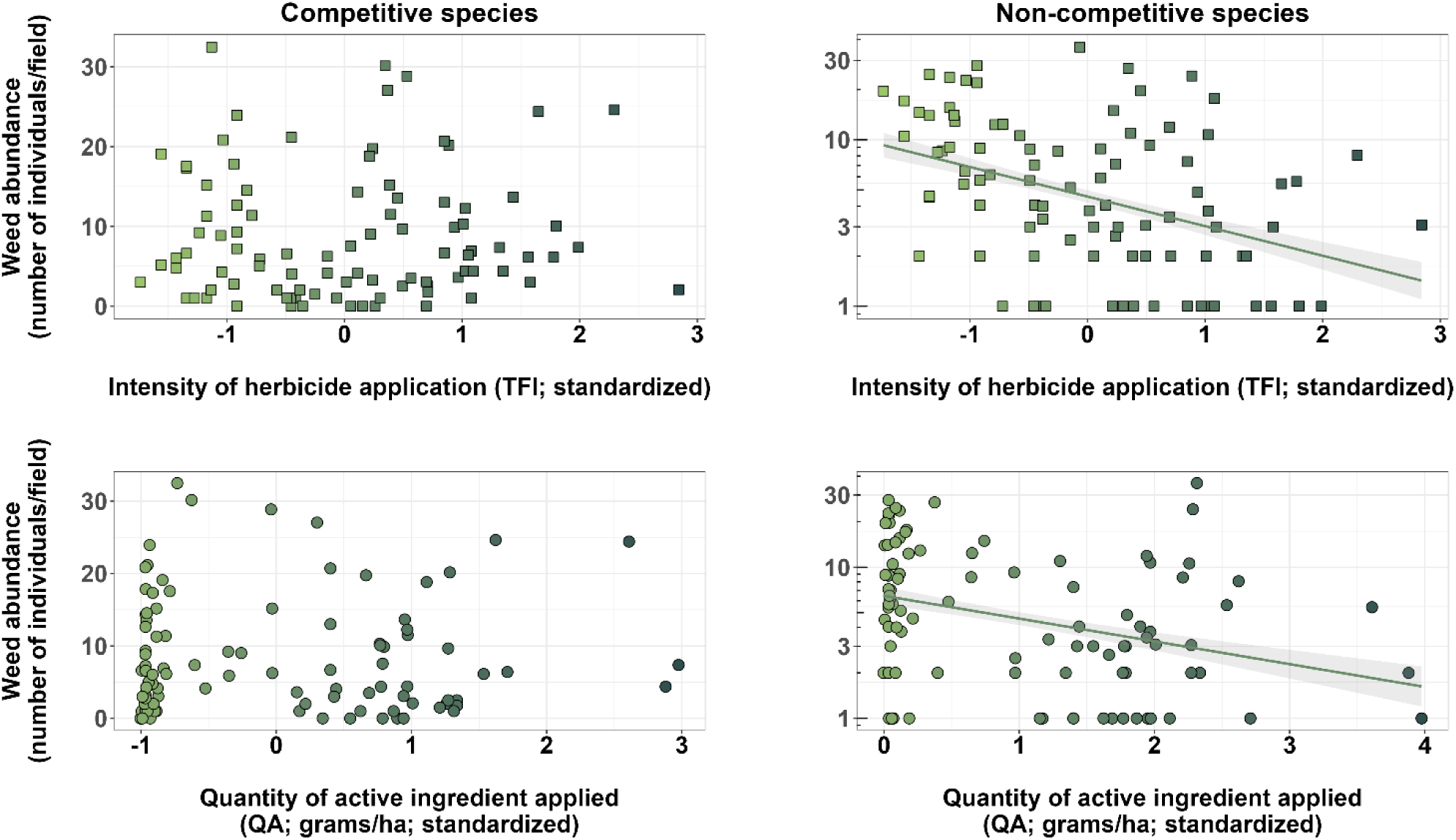
Abundance of competitive and non-competitive weed species in relation to standardized TFI and QA of herbicides. A change of one unit (x-axis) corresponds to an increase or a decrease of one standard deviation from the mean of TFI or QA of herbicides. Linear regression lines are presented when there are evidence supporting a relationship between weed abundance and herbicide input. Grey area represents the 95% confidence interval for the fitted line derived from predictions of the linear mixed-effects model, accounting for the effects of other variables in the model. The partial correlation coefficient, r, indicates the strength and direction of the association between two variables while accounting for other variables in the model. It is displayed for each association. A log scale is used on the y-axis to reflect that the response variable was log-transformed in the model. To avoid issues with log(0) for zero abundance values, we added 1 to all values prior to log-transformation. As a result, a value of 1 on the log-transformed scale corresponds to an observed abundance of 0.

**Table 3:**
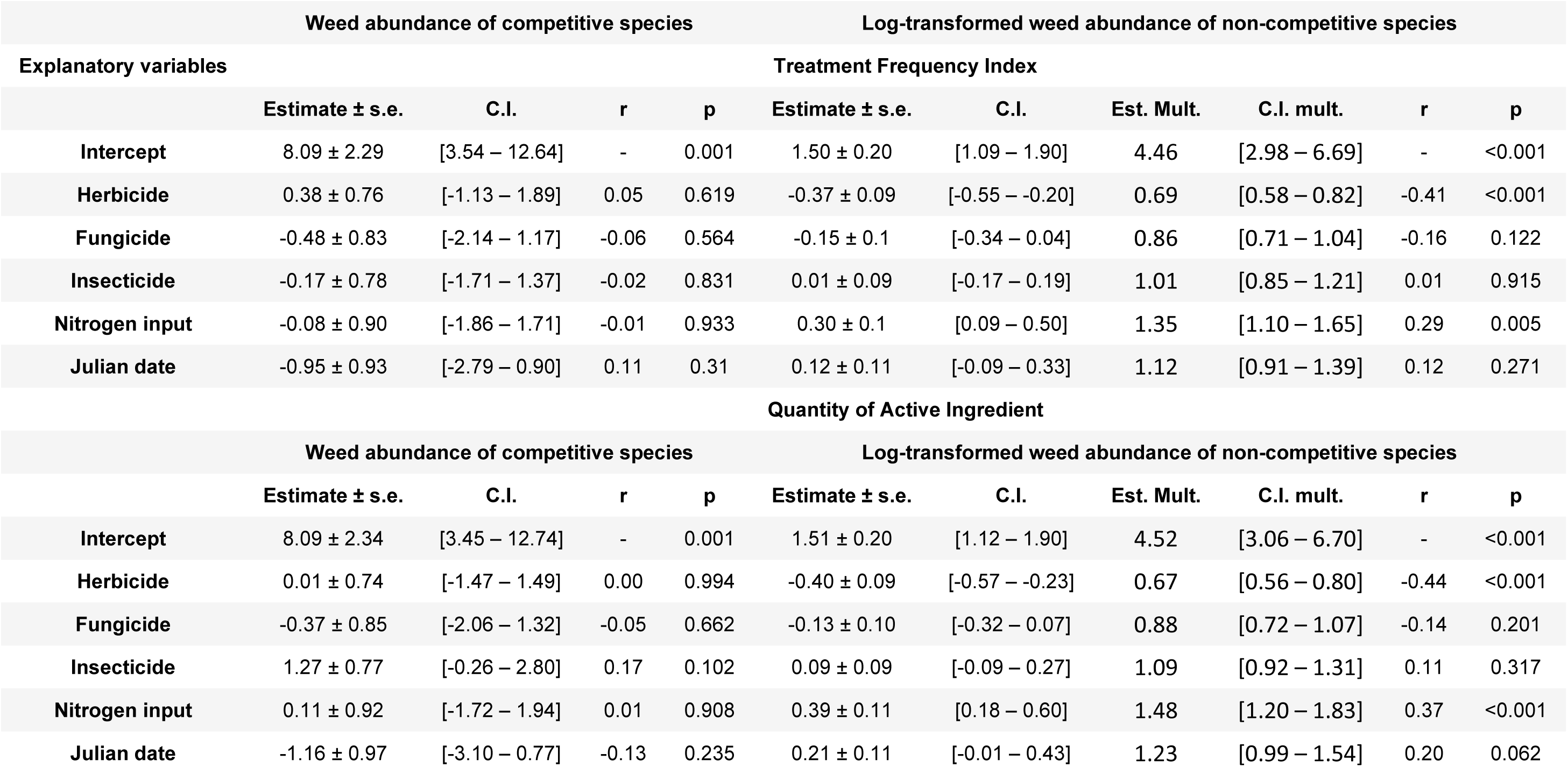
Summary of the linear mixed model investigating the effect of Treatment Frequency Index and Quantity of Active Ingredient of herbicide, fungicide, insecticide, nitrogen input and Julian date (all variables standardized) on weed abundance of competitive and non-competitive species using a restricted maximum likelihood function. The table shows model estimates ± standard error (Est. ± s.e.) and associated 95% confidence intervals (C.I.). P-values are derived from a t-test testing against the null hypothesis that the estimate is 0. The table also shows multiplicative estimates for untransformed variable (Est. mult.) and associated multiplicative confidence intervals (C.I. mult.). Additionally, effect sizes *r* is provided, along with p-values that are derived from a t-test testing against the null hypothesis that the estimate is 0.

### 2.4. Nestedness versus turn-over in beta-diversity response patterns

Since both herbicides and nitrogen influence weed communities, we further analysed β-diversity patterns in response to increasing herbicide while controlling for nitrogen input. Results indicate that TFI of herbicide does not explain total dissimilarity between pairwise fields (βtotal; partial Mantel statistic r = 0.02, p = 0.32), nor does it account for turnover (βBal; r = 0.02, p = 0.36) or nestedness (βGra; r = -0.001, p = 0.49). In contrast, the QA of herbicides affect both turnover (βBal; r = 0.12, p = 0.003) and total dissimilarity (βtotal; r = 0.12, p = 0.005), but not nestedness (βGra; r = -0.06, p = 0.96).

We do not find any difference in beta-diversity in field margin communities, regardless of whether TFI or QA of herbicides is used (all r < 0.07, p > 0.10).

## 3. Discussion

In this study, we investigated the impact of varying levels of herbicide, fungicide, and insecticide use on weed communities across 96 integrated and conventionally managed cereal fields. Our findings reveal that both the intensity and quantity of herbicide applications influence changes in weed communities, with quantity having a stronger impact than intensity on all parameters studied. We also find that the intensity of fungicide applications appears to reduce both weed abundance and species richness, whereas the quantity applied does not. Lastly, insecticides are found to be unrelated to any of the parameters describing weed communities.

Concerning insecticides, a floor effect due to their presence in only a small portion of the studied fields may explain the lack of notable impacts on weed communities. Additionally, their mode of application and the type of data collected in this study may also contribute to blurring any potential effect of insecticides on weeds. Indeed, our study only include data about direct aerial applications, although some products, particularly until 2018, could be applied as seed coating. Therefore, we lack an important piece of information about the intensity and quantity of insecticides applied.

The fact that the intensity of application of fungicides, i.e. higher or lower doses relative to the recommendations, rather than the quantity applied decreased abundance and species richness of weeds suggests that fungicide intensity of application, which may reflect low-dose but frequent applications, could exert indirect and long-term effects on weed suppression and highlights the necessity to reduce the application intensity and not exceed the recommended doses. Such an effect of fungicides may be caused by an alteration of symbiotic arbuscular mycorrhizal fungi in the soil and interference with nutrient uptake by the plants, which may allow certain weed species to dominate, thus resulting in lower overall species richness (Bennett and Cahill Jr, 2016; Francis and Read, 1995; Han et al., 2021). For example, in nutrient-poor dry grasslands, the suppression of arbuscular mycorrhizal fungi through fungicide application has been shown to significantly alter plant species composition, leading to decreased richness and increased dominance by specific grass species (Dostálek et al., 2013). This suggests that fungicides can indirectly shape weed community dynamics by disrupting mutualistic fungal associations. In addition, frequent fungicide applications – particularly during early growth stages – may also exert direct effects on plant growth by inhibiting through inhibition of regulators enzymes (Nordmeyer and Koch, 2014).

We find an impact of both TFI and QA of herbicide on weed community composition through the multivariate analyses. We acknowledge that the variables included in our models explain only a modest proportion of the variance, which is common in community ecology due to the influence of numerous unmeasured or stochastic factors (Greenacre and Primicerio, 2014). However, we did not find any clusters of weed communities, suggesting that communities gradually change as herbicide use increases. This effect may occur through herbicides selectively eliminating more sensitive, non-targeted weed species while allowing others to proliferate (Gaba et al., 2016). In fact, although not rare, weed species that we classified as non-competitive generally do not compete with crops even under low herbicide input (apart from *Chenopodium album* that competes with other crops than winter wheat) and therefore are unlikely to be responsible for yield loss (https://wikiagri.fr/). In contrast, species found to be associated with high herbicide input are known to be competitive weeds. Competitive weeds compete directly with crops, including cereals, for light, nutrients, and water, but may also release allelopathic toxins that hinder crop growth, act as vectors for diseases and therefore impact harvest quality (e.g., increasing moisture in harvested grains) and reduce yields (Tilman, 2002, Wikiagri, 2024, Zimdahl, 2018, Zimdahl, 2008). While farmers may adjust herbicide application in response to the presence of competitive weeds, our models examining the relationship between herbicide input and weed abundance in competitive and non-competitive weeds indicate that greater herbicide intensity and quantity is associated with a lower abundance of non-competitive species, but is unrelated to the abundance of competitive weeds – the primary targets of herbicide. This finding challenges the hypothesis that higher herbicide input simply reflects a response to higher competitive weed pressure. The lack of relationship between competitive species and the intensity and quantity of herbicide further hints at their potential tolerance to herbicides and/or their ability to thrive in environments where competitors have been suppressed by herbicides. Overall, it appears that higher herbicide application gradually eliminates weeds, disproportionately impacting weed species with low competitiveness against crops while favouring those that are more tolerant or resistant to herbicide, and/or are highly competitive against crops (Andert et al., 2022; Gaba et al., 2016; Storkey et al., 2011). The persistence of these problematic species under high herbicide input may be attributed to the evolution of herbicide resistance mechanisms. Herbicide resistance is a well-recognized issue in modern agriculture, particularly among competitive weeds like *Lolium multiflorum* and *Alopecurus myosuroides*, both associated with high herbicide use in our study. These species have evolved resistance to multiple herbicides through both target-site mechanisms, such as mutations or upregulated expression of genes encoding the herbicide’s target enzyme, and non-target-site mechanisms, which involve physiological adaptations like enhanced herbicide metabolism (Brunharo and Tranel, 2023; Franco-Ortega et al., 2021; Gaines et al., 2020).

While herbicide application is the major factor shaping weed communities, our findings also indicate that nitrogen input influences community composition, though to a lesser extent than herbicides, by increasing weed abundance without reducing species richness. Indeed, nitrogen levels are known to modify species composition without decreasing species richness (Humann-Guilleminot et al., 2023; Jiang et al., 2018; Kordbacheh et al., 2023). Several weed species are known to thrive in nitrogen-rich environments, where they outcompete other species, leading to shifts in the overall community composition (Berger et al., 2007; Kim et al., 2006; Moreau et al., 2014). Our results also confirm that nitrogen is a weaker driver of beta diversity than herbicides (Carrié et al., 2022).

We can think of several mechanisms to explain the observed changes in weed communities. Changes in species community composition can occur through species replacement (captured by the balanced variation of β-diversity, i.e. species turnover), shifts in the abundance of pre-existing species (represented by gradient variation of β-diversity, i.e. nestedness), or a combination of both (reflected in total β-diversity dissimilarity). We do not find any relationship between any indices of β-diversity and the intensity of herbicide application, suggesting that TFI only reduces species richness without altering dissimilarity across communities. In contrast, the quantity of herbicide applied appears to notably influence species turnover, overall β-diversity dissimilarity, weed abundance and species richness. This pattern describes both a decline in non-competitive species and their replacement by persistent, more abundant, competitive species while herbicide increases.

The role of weed species in agricultural fields as an essential component of biodiversity is now widely recognized (Marshall et al., 2003; Petit et al., 2011). Numerous studies have highlighted agricultural intensification – comprising herbicide application – as the primary driver behind changes in species richness, overall abundance, and the prominence of certain species (Carmona et al., 2020; Fonderflick et al., 2020; Fried et al., 2010, 2008; Storkey et al., 2011). However, alternative agroecological approaches aim to promote practices that support more environmentally friendly production, contribute to climate change mitigation and adaptation, and minimize or eliminate the use of chemical inputs (Aguilera et al., 2020; Gliessman, 2022; Pimbert, 2015). These strategies include landscape and farm diversification, intercropping, crop and pasture rotation, and the use of biological pest control (Beillouin et al., 2019; HLPE, 2019; Wezel et al., 2020). In addition, agroecological practices encourage diverse weed communities, which serve as an insurance policy that enhances the resilience and agroecosystems services (Wezel et al., 2014; Wojtkowski, 2006; Yachi and Loreau, 1999). In contrast, a reduction in weed diversity could lead to a narrower range of these services, with the residual functions being mainly fulfilled by species that withstand heavy herbicide use and tend to have similar roles (Loreau et al., 2021). Agroecological practices have been shown to promote ecosystem services across different trophic levels, from enhancing soil microbial biomass to increasing pollinator abundance and diversity (Bezner Kerr et al., 2023; Dangles and Casas, 2019; Sandén et al., 2018). Weed diversity plays a key role in supporting these services, not only by contributing to overall biodiversity, but also by attracting beneficial insects, supporting pest regulation, and stabilizing soils (Bretagnolle and Gaba, 2015; Petit et al., 2011). Yet, under conventional farming systems, weeds are still considered as pests species that compete with crops for water, light, and nutrients, and serve as hosts for pests and diseases, ultimately leading to yield reductions (Gazoulis et al., 2024).

This study, which focuses on the impact of pesticides in non-organic cereals fields, reveals how the quantity, and to a lesser extent intensity, of herbicides applied alters weeds communities at all levels from weed species richness to community composition. In particular, herbicides decrease the abundance of non-competitive weeds while replacing them by competitive to crops species. Furthermore, our results indicate that the intensity of application of fungicide also decrease weed abundance and species richness in cereals fields. We acknowledge that this study is limited to cereal crops and similar analyses across other cropping systems – especially those with higher pesticide inputs – are needed to assess the broader applicability of our findings. Furthermore, our study is observational and does not allow for direct causal inferences; experimental approaches would be necessary to confirm the causal effects of herbicide use on competitive and non-competitive weeds. Nonetheless, our study relies on a large sample size and realistic, up-to-date, agricultural practices. Importantly, our results point at a potential limitation of the use of herbicides to control highly competitive weed species and the need to limit their application in favour of agroecological practices that promote the diversity of weeds beneficial to agroecosystems.

## 4. Material and methods

### 4.1. Study area

The study was conducted between 2017 and 2020 in the Long-Term Socio-Ecological Research (LTSER) platform “Zone Atelier Plaine & Val de Sèvre” (hereafter referred to as ZA PVS, Bretagnolle et al., 2018a). The study focuses on integrated and conventional winter cereals fields, which account for ca. 60% of the cultivated crops in the study site, with an average field size of 4.5 ha. The study area encompasses 435 km^2^ of agricultural land located in the southern region of the city of Niort, within the Deux-Sèvres department in France.

### 4.2. Farmers’ survey

Information on farming practices, including pesticide and fertilizer applications (product names, dates and quantities of each application), tillage techniques and the mechanical weed control operations (number, deepness), were obtained through farmer’s interviews performed at the end of each cropping season. Each field in which relevés and interviews were performed were randomly selected before collecting the data. More importantly, the persons in charge of doing the vegetation relevés were blind to the amount of pesticides applied in the fields. Based on the collected information, we derived two widely used quantitative pesticides indicators and nitrogen input values (see Catarino et al., 2019 for details on calculation). The first is the Treatment Frequency Index (TFI) that is a measure of pesticide use intensity, i.e. the number of registered doses applied, for each pesticide, per hectare and per crop season, calculated as follow: 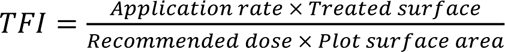. The second is the Quantity of Active Ingredients applied (QA), as the sum of the amount, in grams, of each active ingredient applied per hectare (Möhring et al., 2019). The calculation and use of both indices are thoroughly explained in Möhring et al. (2019), with a clear illustration provided in their Figure 1.

### 4.3. Weed surveys

Identity and occurrence of weeds were recorded each year from April to July in a total of 96 different winter cereals fields, including winter wheat and winter barley (see Bretagnolle et al., 2018b for general methods). Within each field, 20 quadrats (1m² each) - each subdivided into four 0.25 m^2^ subquadrats - are positioned along two parallel transects from field border to centre (10 per transect). Additionally, 5 quadrats (again, subdivided into four 0.25 m^2^) located at one of the field margins are also surveyed (refer to Figure S1 in Bretagnolle et al., 2018b for more details). Number of quadrats and their size were selected to optimally represent the plant community while minimizing survey time (Bretagnolle et al., 2018b). Weed species and abundance were recorded by direct counting with a presence/absence scale within each 0.25 m^2^ sub-quadrat of each plant present. The abundance scale was defined as follows: 0 for absence; 1 when a single individual of a species was present; and 2 if a species was represented by more than one individual. Weed abundance was first calculated at 1m² quadrat, as the sum of abundances of all plants per species present in the four subquadrats, taking the geometric mean (3.33) when abundance was coded as 2 in a sub-quadrat (thus, in a quadrat, for a species, abundance varied from 0 to 13.33). Weed abundance was then defined and calculated as the average between the 20 1m² quadrats in the centre of the field and as the average between the five 1m^2^ quadrats at the field margins.

### 4.4. Statistical analyses

All statistical analyses were performed using the R v. 4.2.2 software (R Core Team, 2022). They were performed separately between field cores and field margins, acknowledging that these distinct areas harbour different weed communities due to edge and landscape effects, as well as differential exposure to pesticides (Bourgeois et al., 2019; Carrié et al., 2022). Our primary focus was to investigate the impact of pesticides on weed communities, thus only data collected in the centre of each field are presented in the main part of this paper. Nonetheless, results obtained from field margins are also given and used for contextualization (full results are presented in the supplementary material).

All analyses investigating the impact of TFI and QA for all pesticides were conducted separately for each index due to their strong correlation and the sensitivity of the results to small changes in the data (all R^2^ > 0.5, all p < 2e^-16^).

#### 4.4.1. Influence of pesticide application on total weed abundance and species richness

We first examined whether the TFI and QA for herbicides, fungicides, and insecticides influence weed abundance and species richness. For this purpose, we ran four linear mixed-effects models (LMMs), with log-transformed weed abundance and log-transformed species richness as the response variables. The explanatory fixed factors included TFI or QA for herbicides, fungicides, and insecticides, as well as nitrogen input and the Julian date (continuous) of the botanical survey. Year was included as a random factor to account for inter-annual variability. All continuous variables were standardized (centered and scaled). LMMs were performed using the *lmer* function from the R package *lmerTest* (Kuznetsova et al., 2017).

#### 4.4.2. Influence of pesticide application at community-level

We addressed the effects of both TFI and QA of herbicides, fungicides and insecticides at the community level. To do this, we used partial distance-based redundancy analyses (partial dbRDA) – an extension of dbRDA (Legendre and Anderson, 1999) – models including either TFI or QA of herbicides, fungicides and insecticides, nitrogen input and Julian date as explanatory fixed factors using *dbrda* function. As a constrained ordination method, dbRDA directly relates species composition to environmental variables, making it well-suited for hypothesis-driven studies. Unlike unconstrained methods (e.g., NMDS, PCA), which do not incorporate explanatory variables, dbRDA are reliable for analysing species–environment relations. It also accommodates the non-Euclidean and zero-inflated nature of species abundance data by allowing the use of suitable dissimilarity measures. Compared to canonical correspondence analysis (CCA), which assumes unimodal species responses, dbRDA assumes linear relationships, making it more suitable for analysing moderate-length gradients like our pesticide gradient (Jupke and Schäfer, 2020). We controlled for the sampling year by adding the year as a conditional variable to the model to account for the variation associated with it. Weed communities were characterized by a response matrix, which contained dissimilarity values across the sites (fields). All explanatory variables were standardized (centered and scaled). We applied a Hellinger transformation on the weed community matrix, i.e. it converts species abundances into the square-root of relative abundance within each site, to better linearize the distances among sites and reduce the effect of dominant species using the *decostand* function (Legendre and Legendre, 2012). Dissimilarity index selection was based on a Spearman rank correlation among seven dissimilarity indices, using the *rankindex* function. This function computes the distance indices between all pairs of samples and correlates these distances with the environmental variables included in our final model using the Spearman correlation method. The purpose was to find a distance measure that accurately represents the original distances in reduced-dimensional space. Among the tested distance, Canberra distance was identified as the most suitable for our data since it is adapted to heterogeneous scales and positive values. We assessed the significance of each explanatory variables by conducting an ANOVA using the *anova.cca* function with 999 permutations. All the functions used for these analyses were done using the R *vegan* package (Oksanen et al., 2022).

#### 4.4.3. Influence of pesticide application on weed abundance of competitive and non-competitive species

After identifying the species responsible for community changes via dbRDA described in the previous section, we grouped them based on their association with high and low herbicide inputs into competitive and non-competitive species, respectively. We then assessed the relative abundance of these groups, exploring how the Treatment Frequency Index (TFI) and the Quantity of Active Ingredient (QA) of pesticides affected them. To do this, we conducted four linear mixed-effects models (LMMs) using the lmer function from the R package *lmerTest*. For non-competitive species, weed abundance was log-transformed. To avoid issues with log(0) for zero abundance values, we added 1 to all values prior to log-transformation. Our models included TFI or QA for herbicides, fungicides, and insecticides, along with nitrogen input and the continuous Julian date of the botanical survey as fixed factors, and Year as a random factor to adjust for inter-annual variability. All continuous variables were standardized.

#### 4.4.4. Underlying mechanisms under weed community changes

To identify the underlying ecological mechanisms of community change as revealed by the dbRDA, we analysed how weed communities responded to herbicide use by calculating abundance-based dissimilarity indices of β-diversity, split into β-turnover and β-nestedness components using the *beta.pair.abund* function from the *betapart* package (Baselga, 2017). In this methodology, the total dissimilarity between sample pairs (βBC) is divided into balanced variation (βBal) and abundance gradient (βGra). βBal measures the turnover of species abundance between samples, while βGra reflects changes in total abundance between communities, indicating nestedness (Baselga, 2017). To assess the influence of herbicides, quantified either as TFI or QA, on these dissimilarities, we applied partial Mantel tests with the *mantel.partial* function from the *vegan* package. The analysis involves a predictor matrix of Euclidean distances based on TFI or QA differences between fields, and response matrices consisting of the β-diversity dissimilarities (βBC, βBal, βGra). Additionally, we controlled for nitrogen input differences using a third matrix to isolate the specific effect of herbicides. The significance of the relationships is evaluated using Pearson’s coefficient and 999 permutations in the partial Mantel test setup. For measuring dissimilarities, we chose the Bray-Curtis index, determined to be highly suitable for our data by the *rankindex* function, second only to the Canberra index (the latter not being available in the package). This index was applied to a matrix of species abundances across sample sites, transformed using Hellinger to normalize the data.

#### 4.4.5. Modelling assumptions

We chose not to include additional environmental variables or interaction terms between the three pesticides to avoid model complexity and instability (Clark et al., 2020). However, to ensure the validity of this decision, we followed the recommendations of Sutherland et al., 2023 and tested a selection of models with and without additional agricultural practices added alone or in interactions (i.e. mechanical weeding, tillage operations, interactions between pesticides input and nitrogen). The models that included only pesticides, nitrogen inputs and Julian date were consistently selected as the best fit. Modelling assumptions of LMMs (normality and homoscedasticity of residuals, homogeneity of variance, collinearity) were validated by visual inspection using the *check_model* function of the performance R package (Lüdecke et al., 2021). All variance inflation factors were below three.

Multivariate models were validated through a visual inspection of residuals to ensure compliance with assumptions of multivariate normality and homogeneity. This included examining the residual distributions and graphically plotting the residuals against the fitted values.

Following recommendations in the ASA statement on p-values (Wasserstein and Lazar, 2016), we do not interpret our results based on an arbitrary threshold for statistical significance (p < 0.05). Instead, we use p-values to measure the amount of statistical evidence against the null hypothesis and further examine the estimated coefficients of the model to assess the strength of the associations and the confidence we should have in those estimates. Effect sizes, i.e. correlation coefficient (*r*), are provided for each Linear-mixed effect models (LMM) and indicate the strength and direction of the association. The common interpretation is based on Cohen, 1988 guidelines: small (r < 0.30), medium (r < 0.50), large (r > 0.50).

## Data and materials availability

All plant data and R code generated to conduct the analyses described in that study are available in a public database: https://doi.org/10.48579/PRO/9CKQC6. Please note that pesticides and fertilizer data are confidential data under the French data protection law (CNIL). Please contact the authors if you want to obtain this data.

## Acknowledgements

We would like to thank all farmers who participated in the survey for devoting their time and for their confidence expressed by letting us monitor weed flora within their fields. We are deeply grateful to Adrien Berquer, Sébastien Boinot, Valentin Cornet, Stanilas de Closets, Thierry Fanjas-Mercere, Jonathan Millaud and Ségolène Rousselet for their investment in weed flora monitoring. We also would like to thank Ambroise Lahut and Niklas Mörhing for data curation and computation of agricultural practices metrics; and Fabien Vialloux and Claire Lamare for conducting farmers’ interviews. We are grateful to Sylvain Losdat for his advice and validation of our statistical approach.

## Funding

The study has received funding from the CNRS project SeeLife for long-term biodiversity monitoring and the Nouvelle Aquitaine Region (project Transform’ Actions). SG and VB were funded by INRAE and CNRS. SHG received funding from CNRS and an SNSF-Postdoc.Mobility grant (#P500PB_222110).

## Author contributions

**S.H-G. contributed to:** Conceptualization, Methodology, Formal analysis, Data curation, Visualization, Writing - Original Draft, Writing - Review & Editing.

**V.B. contributed to:** Conceptualization, Methodology, Data curation, Supervision, Project administration, Funding acquisition, Writing - Review & Editing.

**S.G. contributed to:** Conceptualization, Methodology, Data curation, Supervision, Project administration, Funding acquisition, Writing - Review & Editing.

## Competing interests

Authors declare that they have no competing interests.

## Notes

https://doi.org/10.48579/PRO/9CKQC6

### Competing Interest Statement

The authors have declared no competing interest.

