## Supplementary tables and figures for "Off target: herbicides applied in cereal fields exclude non-competitive species, while replacing them by competitive weeds"

**Table S1:** Median, mean ± standard deviation (sd), minimum and maximum for species richness and weed abundance per m², Treatment Frequency Index (i.e. intensity of application; TFI) and quantity of active ingredient applied (QA; g/ha) of herbicide, fungicide and insecticide and nitrogen input (g/ha) in 2017, 2018, 2019 and 2020.

| **Year** | | **2017** | **2018** | **2019** | **2020** |
|---|---|---|---|---|---|
| Weed abundance | Median | 11.03 | 41.16 | 27.56 | 26.93 |
| | Mean ± sd | 15.25 ± 10.97 | 46.28 ± 28.11 | 37.97 ± 31.55 | 27.88 ± 17.70 |
| | Min - Max | 3 - 52.46 | 7.5 - 120.02 | 6.16 - 135.16 | 1 - 71.47 |
| Species richness | Median | 8 | 12 | 9.5 | 10 |
| | Mean ± sd | 8.09 ± 4.10 | 11.68 ± 4.71 | 9.82 ± 5.77 | 10.45 ± 5.69 |
| | Min - Max | 2 - 19 | 5 - 24 | 2 - 25 | 1 - 25 |
| TFI of herbicide | Median | 1.76 | 1.73 | 1.80 | 1.35 |
| | Mean ± sd | 1.77 ± 0.72 | 1.69 ± 0.75 | 1.81 ± 0.97 | 1.59 ± 0.74 |
| | Min - Max | 0.5 - 2.94 | 0.6 - 3.1 | 0.5 - 3.93 | 0.36 - 3.12 |
| TFI of fungicide | Median | 1.55 | 1.37 | 1.05 | 0.86 |
| | Mean ± sd | 1.55 ± 0.69 | 1.6 ± 0.86 | 1.14 ± 0.80 | 0.99 ± 0.58 |
| | Min - Max | 0.35 - 2.75 | 0.44 - 3.5 | 0 - 2.6 | 0 - 2.36 |
| TFI of insecticide | Median | 0 | 0 | 0 | 0 |
| | Mean ± sd | 0.32 ± 0.44 | 0.31 ± 0.66 | 0.39 ± 0.55 | 0.27 ± 0.44 |
| | Min - Max | 0 - 1.1 | 0 - 2 | 0 - 2.09 | 0 - 1 |
| QA of herbicide | Median | 1278.5 | 823.75 | 430.76 | 896.96 |
| | Mean ± sd | 1115.38 ± 1002.54 | 1159.53 ± 1146.32 | 999.64 ± 1191.88 | 1205.94 ± 1245.24 |
| | Min - Max | 39.85 - 3000 | 10.6 - 3994.47 | 15 - 4400 | 0 - 4294.3 |
| QA of fungicide | Median | 742.5 | 587.75 | 431.25 | 291.5 |
| | Mean ± sd | 613.71 ± 273.62 | 822.85 ± 759.47 | 567.25 ± 748.47 | 416.86 ± 565.05 |
| | Min - Max | 120 - 1050 | 150 - 2933.34 | 0 - 2776.7 | 0 - 2658.7 |
| QA of insecticide | Median | 0 | 0 | 0 | 0 |
| | Mean ± sd | 5.71 ± 9.06 | 6.53 ± 14.80 | 10.53 ± 26.64 | 5.56 ± 10.12 |
| | Min - Max | 0 - 27.5 | 0 - 50 | 0 - 97.36 | 0 - 25 |
| Nitrogen input | Median | 167.70 | 184.10 | 20.00 | 157.95 |
| | Mean ± sd | 163.40 ± 47.32 | 174.49 ± 43.90 | 76.97 ± 81.82 | 141.08 ± 76.45 |
| | Min - Max | 0 – 225.40 | 0 – 260.13 | 0 – 197.90 | 0 – 239.60 |

**Table S2:** Summary of the linear mixed model investigating the effect of Treatment Frequency Index and Quantity of Active Ingredient of herbicide, fungicide, insecticide, nitrogen input and Julian date (all variables standardized) on log-transformed weed abundance and log-transformed species richness in field margins using a restricted maximum likelihood function. The table shows model estimates ± standard error (Est. ± s.e.) and associated 95% confidence intervals (C.I.). The table also shows multiplicative estimates for untransformed variable (Est. mult.) and associated multiplicative confidence intervals (C.I. mult.). Additionally, the effect size r is provided, along with p-values that are derived from a t-test testing against the null hypothesis that the estimate is 0.

| | **Log-transformed weed abundance** | | | | | | **Log-transformed species richness** | | | | | |
|---|---|---|---|---|---|---|---|---|---|---|---|---|
| **Explanatory variables** | **Treatment Frequency Index** | | | | | | | | | | | |
| | **Estimate ± s.e.** | **C.I.** | **Est. Mult.** | **C.I. mult.** | ***r*** | **p** | **Estimate ± s.e.** | **C.I.** | **Est. Mult.** | **C.I. mult.** | ***r*** | **p** |
| **Intercept** | 3.61 ± 0.2 | [3.21 – 4.01] | 36.92 | [24.67 – 55.26] | - | <0.001 | 2.46 ± 0.07 | [2.31 – 2.60] | 11.66 | [10.12 – 13.43] | - | <0.001 |
| **Herbicide** | -0.1 ± 0.08 | [-0.25 – 0.05] | 0.91 | [0.78 – 1.05] | -0.14 | 0.201 | -0.08 ± 0.07 | [-0.22 – 0.07] | 0.92 | [0.80 – 1.07] | -0.11 | 0.286 |
| **Fungicide** | -0.07 ± 0.08 | [-0.23 – 0.09] | 0.93 | [0.80 – 1.09] | -0.09 | 0.388 | 0 ± 0.08 | [-0.15 – 0.15] | 1 | [0.86 – 1.16] | 0 | 0.995 |
| **Insecticide** | 0.03 ± 0.07 | [-0.12 – 0.17] | 1.03 | [0.88 – 1.19] | 0.04 | 0.734 | 0.02 ± 0.07 | [-0.13 – 0.16] | 1.02 | [0.88 – 1.18] | 0.03 | 0.806 |
| **Nitrogen input** | 0.17 ± 0.09 | [-0.01 – 0.34] | 1.18 | [0.99 – 1.41] | 0.2 | 0.06 | 0.22 ± 0.08 | [0.07 – 0.37] | 1.25 | [1.07 – 1.45] | 0.3 | 0.004 |
| **Julian date** | 0.11 ± 0.09 | [-0.07 – 0.29] | 1.11 | [0.93 – 1.33] | 0.13 | 0.228 | 0.14 ± 0.07 | [-0.01 – 0.28] | 1.15 | [0.99 – 1.33] | 0.2 | 0.06 |
| | **Quantity of Active Ingredient** | | | | | | | | | | | |
| | **Log-transformed weed abundance** | | | | | | **Log-transformed species richness** | | | | | |
| | **Estimate ± s.e.** | **C.I.** | **Est. Mult.** | **C.I. mult.** | ***r*** | **p** | **Estimate ± s.e.** | **C.I.** | **Est. Mult.** | **C.I. mult.** | ***r*** | **p** |
| **Intercept** | 3.61 ± 0.21 | [3.20 – 4.02] | 36.95 | [24.50 – 55.74] | - | <0.001 | 2.46 ± 0.07 | [2.32 – 2.60] | 11.66 | [10.13 – 13.41] | - | <0.001 |
| **Herbicide** | -0.07 ± 0.08 | [-0.22 – 0.08] | 0.94 | [0.81 – 1.09] | -0.09 | 0.382 | -0.05 ± 0.07 | [-0.20 – 0.09] | 0.95 | [0.82 – 1.10] | -0.07 | 0.491 |
| **Fungicide** | -0.01 ± 0.08 | [-0.17 – 0.15] | 0.99 | [0.84 – 1.16] | -0.01 | 0.896 | -0.03 ± 0.08 | [-0.18 – 0.13] | 0.97 | [0.83 – 1.14] | -0.04 | 0.738 |
| **Insecticide** | 0.1 ± 0.07 | [-0.05 – 0.25] | 1.1 | [0.95 – 1.28] | 0.14 | 0.185 | 0.1 ± 0.07 | [-0.04 – 0.24] | 1.1 | [0.96 – 1.27] | 0.14 | 0.173 |
| **Nitrogen input** | 0.18 ± 0.09 | [-0.00 – 0.36] | 1.2 | [1.00 – 1.43] | 0.21 | 0.053 | 0.26 ± 0.08 | [0.10 – 0.41] | 1.29 | [1.11 – 1.50] | 0.33 | 0.001 |
| **Julian date** | 0.11 ± 0.09 | [-0.08 – 0.30] | 1.12 | [0.92 – 1.34] | 0.12 | 0.251 | 0.14 ± 0.08 | [-0.01 – 0.30] | 1.15 | [0.99 – 1.35] | 0.19 | 0.066 |

0

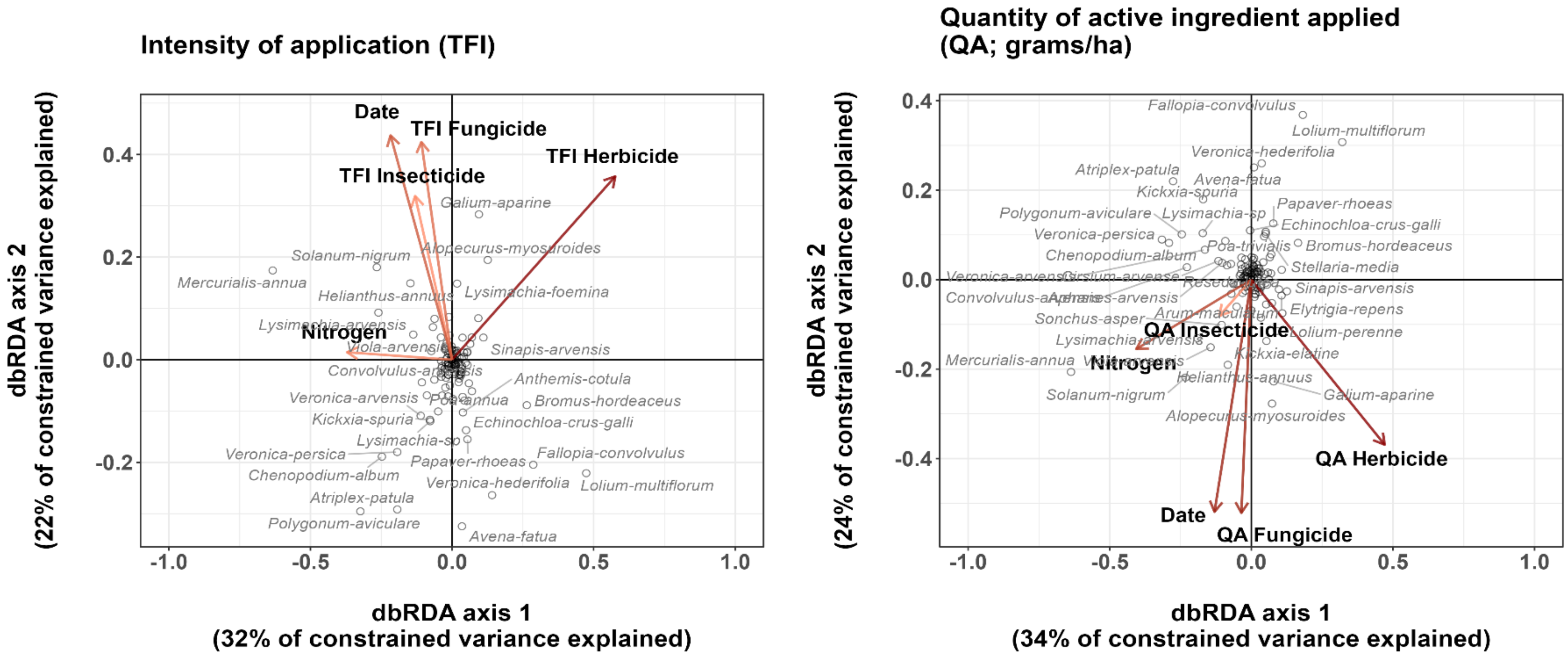


**Figure S1:** dbRDA biplot projections displaying weed community composition along herbicide use gradients based on the intensity of application of herbicide (Treatment Frequency Index, TFI; left) and the quantity of active ingredient applied (QA; right) in field centres. In both models, herbicide input contributes most strongly to the first dbRDA axis (dbRDA1). The models explain 6.52% (TFI) and 7.08% (QA) of the variance in community composition. The vectors represent the direction and strength of the environmental variables, i.e. the effect size, with longer and redder vectors indicating a stronger correlation with the axes. Dots represent individual plant species, with labels displayed only for weeds that most significantly contribute to the observed patterns (those with scores between -0.1 and 0.1). The axes depict gradients of variation, with dbRDA1 accounting for the largest portion of variance. Scores on dbRDA1 are predominantly associated with herbicide levels (both TFI and QA). Scores on dbRDA2 are mainly linked to fungicide levels and Julian dates.

8

**Table S3:** Summary statistics from the dbRDA model and ANOVA using Canberra distance to test the influence of TFI/QAs of herbicide, fungicide, insecticide, nitrogen inputs, and Julian date on weed community composition in field margins. The table shows the total contribution of each variable to the variance in community composition and the proportion of variance explained by each axis. It also includes the coefficients for each variable on the ordination axes, representing their coordinates along each axe. Additionally, the effect size, indicating the strength of the influence of each variable on the ordination, is provided, along with p-values. Effects sizes correspond to the length of each arrow in the biplot and are calculated as the Euclidean distance from the origin to the variable's coordinates on dbRDA1 and dbRDA2 (Figure S2). All variables were standardized (centered and scaled).

| | **Treatment Frequency Index (total contribution of all variables = 5.76%)** | | | |
|---|---|---|---|---|
| **Explanatory variables** | **Coefficient associated with each explanatory variable on each axis** | | | |
| | **dbRDA1** | **dbRDA2** | **Effect size** | **p** |
| Herbicide | -0.01 | 0.55 | 0.81 | 0.28 |
| Fungicide | -0.22 | -0.14 | 0.38 | 0.70 |
| Insecticide | -0.29 | -0.08 | 0.43 | 0.54 |
| Nitrogen input | -0.14 | -0.67 | 1.00 | 0.17 |
| Julian date | -0.77 | 0.10 | 1.14 | 0.02 |
| Proportion explained of each axe:<br>dbRDA 1 = 26.33% (0.15% of variance explained by dbRDA1 overall)<br>dbRDA 2 = 23.20% (0.13% of variance explained by dbRDA2 overall) | | | | |
| | **Quantity of Active ingredient (total contribution of all variables = 6.19%)** | | | |
| **Explanatory variables** | **Coefficient associated with each explanatory variable on each axis** | | | |
| | **dbRDA1** | **dbRDA2** | **Effect size** | **p** |
| Herbicide | 0.29 | 0.75 | 1.17 | 0.05 |
| Fungicide | 0.31 | -0.36 | 0.64 | 0.17 |
| Insecticide | 0.43 | -0.18 | 0.63 | 0.52 |
| Nitrogen input | 0.17 | -0.39 | 0.65 | 0.08 |
| Julian date | 0.72 | 0.03 | 1.04 | 0.04 |
| Proportion explained of each axe:<br>dbRDA 1 = 25.58% (0.14% of variance explained by dbRDA1 overall)<br>dbRDA 2 = 24.70% (0.13% of variance explained by dbRDA2 overall) | | | | |

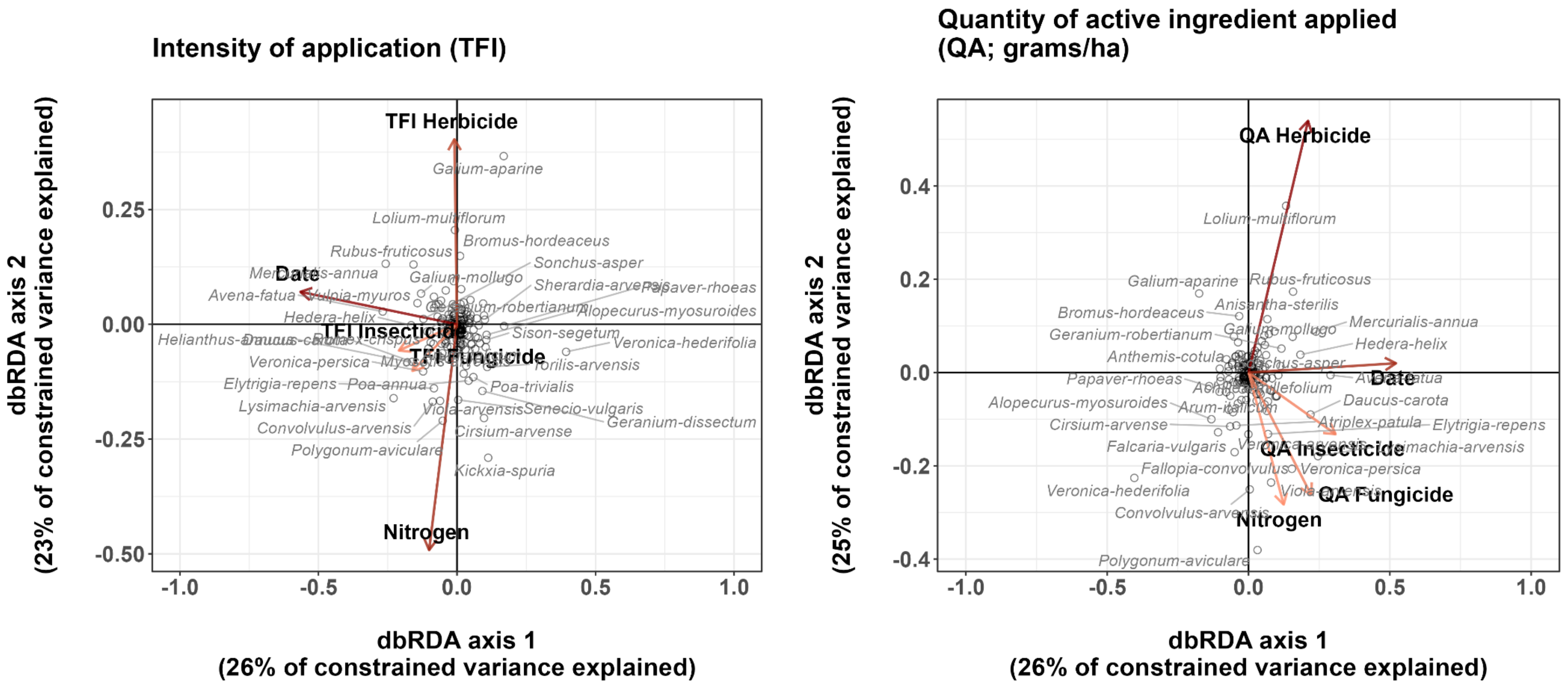


**Figure S2:** dbRDA biplot projections displaying weed community composition along herbicide use gradients based on the intensity of application of herbicide (Treatment Frequency Index, TFI; left) and the quantity of active ingredient applied (QA; right) in field margins. In both models, herbicide input contributes most strongly to the first dbRDA axis (dbRDA1). The models explain 5.76% (TFI) and 6.19% (QA) of the variance in community composition. The vectors represent the direction and strength of the environmental variables, i.e. the effect size, with longer and redder vectors indicating a stronger correlation with the axes. Dots represent individual plant species, with labels displayed only for weeds that most significantly contribute to the observed patterns (those with scores between -0.1 and 0.1). The axes depict gradients of variation, with dbRDA1 accounting for the largest portion of variance. Scores on dbRDA1 are predominantly associated with herbicide levels (both TFI and QA). Scores on dbRDA2 are mainly linked to fungicide levels and Julian dates.
